# C-Eci: A Cubic-Eci Combined Clearing Method For 3D Follicular Content Analysis In The Fish Ovary

**DOI:** 10.1101/2020.03.05.978189

**Authors:** Manon Lesage, Manon Thomas, Jérôme Bugeon, Adèle Branthonne, Stéphanie Gay, Emilie Cardona, Julien Bobe, Violette Thermes

## Abstract

Deciphering mechanisms of oocyte development in female fishes still remains challenging and a comprehensive overview of this process at the level of the organ is still needed. The recent development optical tissue clearing methods have tremendously boosted the 3D imaging of large size biological samples that are naturally opaque. However, no attempt of clearing on fish ovary that accumulates extremely high concentration of lipids within oocytes has been reported to date. To face with this ovarian-specific challenge, we combined two existing clearing methods, the non-toxic solvent-based Eci method for efficient clearing and the CUBIC method to enhance lipid removal and reduce non-specific staining. The methyl green fluorescent dye was used to stain nuclei and delineate follicles. Using this procedure (named C-Eci), ovaries of both medaka and trout could be imaged in 3D and all follicles analyzed. To our knowledge this is the first procedure elaborated for clearing and imaging fish ovary in 3D. The C-Eci methods thus provides an interesting tool for getting precise quantitative data on follicular content in fish ovary and promises to be useful for further morphological studies.

**Summary:** A modified ethyl-cinnamate-based clearing method allows solving fish ovary-specific challenges for 3D imaging, including high lipid-contents, and analyzing the ovarian follicular content in medaka and trout.

## INTRODUCTION

Although much effort has been made in recent years towards understanding oogenesis mechanisms at a cellular level in fish, we still lack a comprehensive overview of the process at the level of the organ. Tightly regulated networks of hormones, secreted factors and intrinsic signaling pathways underlies the progression of each single oocyte throughout oogenesis(Lubzens et al., 2010)(Nakamura et al., 2010). In terms of timing, it however exists different dynamics of oogenesis in adult females, in relation with the different reproductive strategies that exist in fish(McBride et al., 2015). So far, studies related to oogenesis in fish have mainly been based on follicular (*i.e.* oocytes and their surrounding somatic supporting cells) content analyses from ovarian 2D histological sections, which require extensive extrapolations of data (Brown-Peterson et al., 2011). Three-dimensional (3D) analyses of ovarian follicular contents would thus be a major milestone on the path towards a better comprehension of the oogenesis temporal dynamics in fish.

Techniques for 3D imaging of large biological samples that are naturally opaque have experienced major technical breakthroughs this past decade, including the development of numerous optical clearing methods to enhance tissue transparency and reduce light scattering(Richardson and Lichtman, 2015a). These methods are either aqueous-based or solvent-based methods and are grouped into four main classes, including the simple immersion methods, the hyperhydrating methods, the solvent-based methods and the tissue transformation methods(Richardson and Lichtman, 2015b; Silvestri et al., 2016). Simple immersion methods rely on the homogenization of refractive indexes of medium and tissues by using high-refractive index (hRI) aqueous solutions. These solutions are composed of the contrast agent iohexol (also called Histodenz), such as for the RI Matching Solutions (RIMS), or are sugar-based methods, such as for SeeDeepBrain (SeeDB)(Yang et al., 2014a)(Ke et al., 2013)(Affaticati et al., 2017). Methods of the second class are hyperhydrating solutions, such as the Clear Unobstructed Brain/Body Imaging Cocktails and Computational analysis (CUBIC) that is based on the use of the hyperhydrating aminoalcohols, urea and removal of lipids with detergent(Susaki et al., 2014a). Methods of the third class are tissue transformation methods, such as the emblematic PAssive CLARITY Technique (PACT) that use an hydrogel to stabilizes the tissue structure while removing lipids with detergent(Yang et al., 2014b). In the fourth class, the 3DISCO method (3D Imaging of Solvent-Cleared Organs) allows to bypass clearing performance limitations of water-based clearing methods, while preserving fluorescence(Ertürk et al., 2011). More recently, Klingberg *et al.* developed a new clearing method using the ethyl-3-phenylprop-2-enoate (Eci) that is considered nontoxic according to the European directive 67/548/EWG and extremely efficiency for clearing mammalian tissues (Klingberg et al., 2017)(Huang et al., 2019). Some of these methods were already used for clearing the whole mouse ovary and allowed successful 3D imaging, however no attempts on fish ovary have been reported to date(Feng et al., 2017)(Malki et al., 2015a)(Zheng et al., 2014)(McKey et al., 2020).

We tested several of these clearing methods on the medaka ovary and we finally established a protocol combining the CUBIC and Eci methods, which allowed us to solve the fish ovarian-specific challenges and optimize 3D fluorescent imaging of the fish ovary. Using this novel protocol, named C-Eci, ovaries of both medaka and rainbow trout could be cleared and the follicular content analyzed in 3D.

## RESULTS AND DISCUSSSION

### The solvent-based method Eci efficiently clears medaka ovary

We first assessed the clearing efficiency on medaka ovary of various hRI simple immersion solutions, of the hyperhydrating CUBIC method and of solvent-based methods (Eci and iDISCO+). For hRI simple immersion, samples were treated with hRI matching solutions adjusted to increasing refractive index, from 1,353 to 1.49. The apparent transparency in each condition was compared to that of an opaque non-cleared ovary (Fig. 1). With the different hRI solutions, samples displayed only a tiny increase of transparency at the maximum RI (1.49). When treated with CUBIC, transparency was significantly increased. The higher transparency was obtained with the solvent-based Eci and iDISCO+ methods.

**Figure 1:**
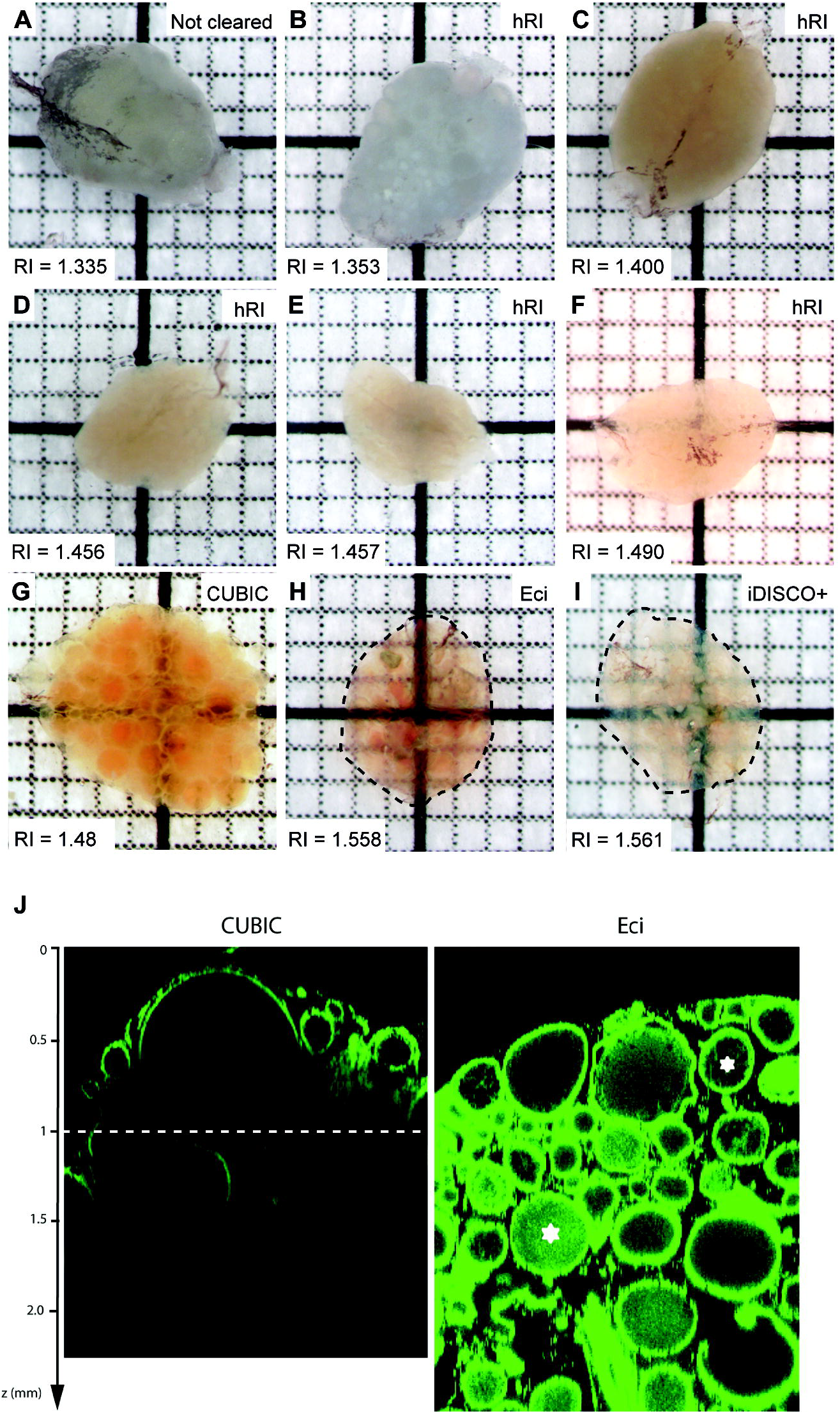
Assessment of the transparency of adult medaka ovaries treated with different clearing methods. Ovaries dissected from adult medaka were cleared with hRI immersion solutions, CUBIC, Eci or iDISCO+ methods. **(A)** Non-cleared ovary in PBS. **(B-F)** Ovaries cleared with hRI immersion solutions with increasing refractive index. **(G-I)** Ovaries cleared with the indicated methods, CUBIC, Eci or iDISCO+. Transparency was assessed by the visualization of black squares. Dotted black line indicates the edge of ovaries after clearing. The solvent-based clearing methods Eci and iDSICO+ are the most efficient methods for clearing medaka ovary. **(J)** Comparison of the fluorescent recovery in depth after CUBIC and Eci methods. The maximal signal recovery in depth was obtained with the Eci method although an important intra-follicular background was observed. Square = 1mm x 1mm.

The optical penetration in depth of confocal imaging after clearing with either CUBIC or Eci methods was then compared. Nuclei, including nuclei of the theca and granulosa cells lining each oocyte, were stained with the Methyl Green (MG) nuclear dye(Prieto et al., 2014). XZ views of z-stacks revealed a maximal signal recovery in depth of less than 1 mm with CUBIC, whereas MG fluorescence was recovered up to 2.5 mm with Eci, corresponding to the limit of the objective working distance (Fig. 1J). However, a nonspecific intra-cytoplasmic background was found for Eci-treated samples, whereas no background was detected in CUBIC-treated samples (Fig. 1J, stars). Similar background was observed in ovaries stained with a less concentrated MG solution (8µg/ml).

These results show that solvent-based methods (iDISCO+ and Eci) enable better clearing of medaka ovary than aqueous-based methods (RIMS, DSP and CUBIC). In mouse, previous studies reported clearing of fetal and adult ovaries with the aqueous-based methods CLARITY(Feng et al., 2017), ScaleA2(Malki et al., 2015b) or the CUBIC method(Kagami et al., 2018). However, higher clearing efficiency of the mouse ovary was also obtained recently by using the solvent-based iDISCO+ method(McKey et al., 2020). In addition, the lipid-content is markedly higher in fish ovary than in mouse ovary, since fish oocytes exhibit higher lipid supply and larger size due to their telolecithal nature. It is thus even more consistent that solvent-based clearing methods, employing the highest RI matching solutions, are more appropriated for clearing fish ovaries than aqueous-based methods. We must however note that a strong intra-follicular background was observed with the Eci method.

### Optimization of the Eci method

Based on the assumption that the intra-follicular background was due to remaining lipids in oocytes after Eci treatment, we tested whether additional lipid removal may improve the staining specificity. Entire ovaries were thus incubated in the aqueous reagent 1 of the CUBIC clearing method (CUBIC-1) prior MG staining and Eci clearing (Fig. 2A)(Susaki et al., 2014b). Furthermore, we tested whether different MG staining conditions may also contribute to reduce the background(Prieto et al., 2015)(Mohtasham et al., 2010)(Sabatani, 1951). CUBIC-1 pre-incubated ovaries (C-Eci protocol) and not pre-incubated ovaries (Eci native protocol) were thus processed for MG staining at two different temperatures (4°C and 50°C) and pH (pH4 and pH7, Fig. 2B). XY-plans at 1000 μm depth from z-stacks of the different cleared-samples were compared. Follicles of different sizes were easily distinguishable in each condition, except with Eci at 50°C-pH4 where almost no fluorescent signal was recovered. Follicles appeared well packed in almost all conditions, except at 4°C-pH4 where follicles appeared distorted and slightly dissociated. The fluorescence ratio signal/background was significantly increased with the C-Eci protocol and the higher ratio was obtained at 50°C-pH7 that offers the best signal contrast (Fig. 2C). To evaluate the effect of clearing on the ovarian size, the apparent surface of ovaries was measured on macroscopic images before and after Eci and C-Eci methods (Fig. 2D). Results show that the apparent ovarian size was significantly reduced after C-Eci treatment (20.51 ± 4.10 mm^2^) compared to non-treated samples (26.01 ± 5.54 mm^2^). Almost no size modification was observed with Eci alone. The shrinkage of ovaries treated with C-Eci were therefore of about 21%.

**Figure 2:**
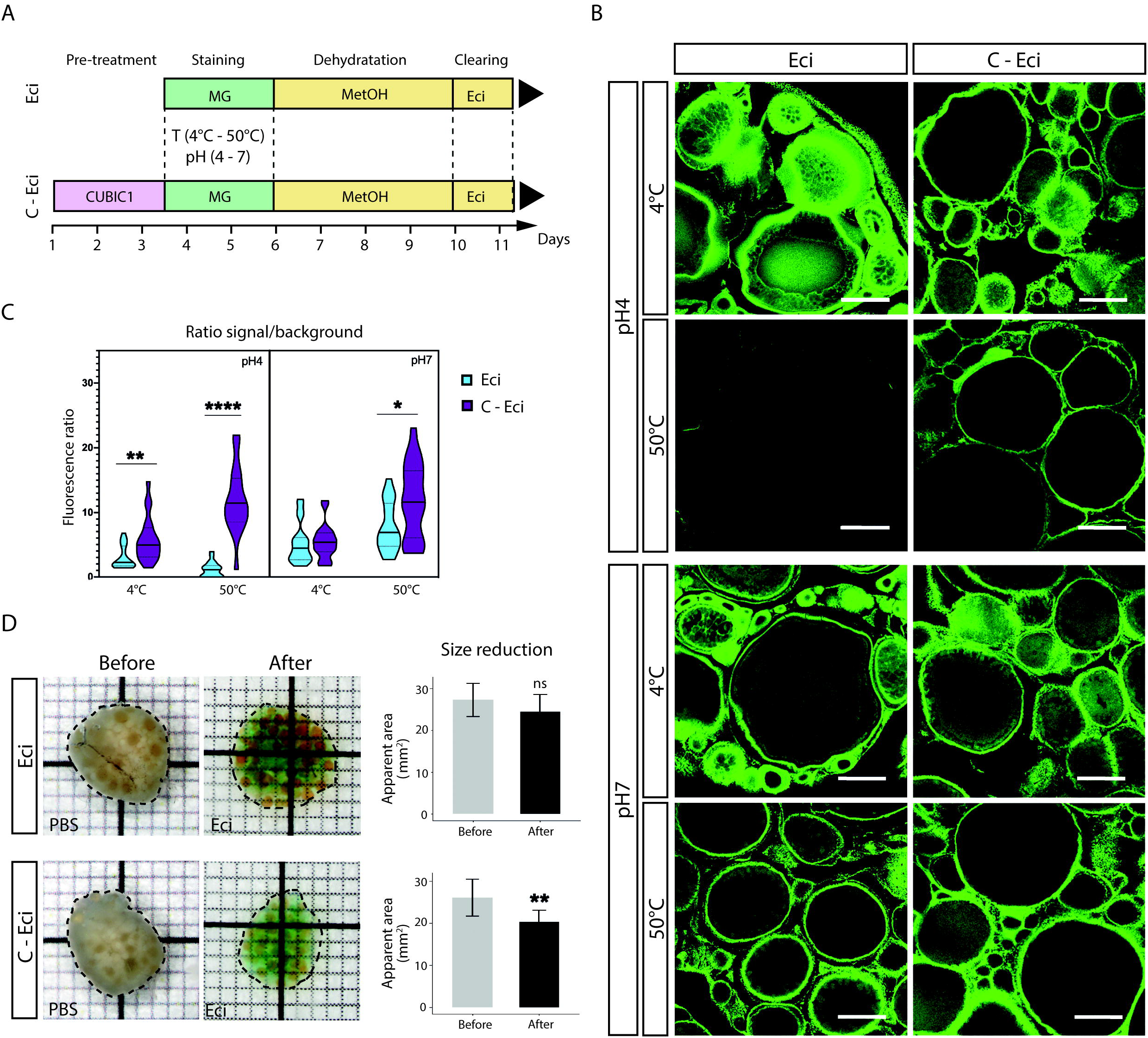
Combining CUBIC and Eci methods efficiently improve nuclear staining of whole medaka ovary. **(A)** Whole-ovary clearing protocol timeline. Medaka ovaries were cleared following the Eci procedure with or without a pretreatment with the aqueous reagent 1 of the CUBIC method (CUBIC-1). Nuclei were stained with methyl green (MG) using different conditions of pH and temperature. **(B)** MG fluorescent signal (in green) acquired from XY planes at 1000 μm depth with the same settings from ovaries treated with Eci or C-Eci. Tissue integrity and intra-cytoplasmic background are improved when samples are treated with C-Eci and when incubated at 50°C for MG staining. **(C)** Quantification of the ratio signal/background of the MG staining. All data are mean ± SD of 20 ROI measured on one ovarian section. **(D)** The apparent ovarian surface areas were measured from brightfield images (black dotted lines) before and after clearing with Eci (n=7) and C-Eci (n=12). The tissue shrinkage is significant with the C-Eci method (Wilcoxon test, p=0.002516). Data are presented as means ± SD. Square = 1 x1 mm; scale bar: 250μm.

These results indicate that combining CUBIC and Eci (C-Eci) significantly reduces intra-follicular background in medaka ovary by increasing delipidation of follicles, and that 50°C-pH7 staining conditions significantly improve the MG staining. We cannot exclude that similar results could also be obtained with the widely used iDISCO+ method, since this later includes a delipidation step with the organic dichloromethane reagent (DCM)(Liebmann et al., 2016)(Renier et al., 2016). However, CUBIC-1 and Eci solutions remain very advantageous since they have a very low toxicity. The C-Eci method is thus an interesting non-toxic alternative to the iDISCO+ method, which greatly facilitates handling samples though similar results might likely be obtained with the iDISCO+ method.

### 3D imaging of a medaka ovary after C-Eci and 3D analysis of follicular content

An ovary collected from an adult medaka female was stained with MG at 50°C-pH7, cleared with the C-Eci protocol and imaged by confocal microscopy (Fig. 3). The whole volume was reconstructed (Fig. 3A). All follicles were detectable and no loss of resolution was observed throughout the sample as shown on XY-planes at different depth (Fig. 3B, top panels). Ovarian follicles were segmented from the 3D reconstruction data by using a computational semi-automatic 3D segmentation procedure (Fig. 3B, bottom panels). The resulting 3D volume reconstruction displays populations of growing follicles of various sizes (Fig. 3C). As mentioned above, the C-Eci treatment induces a global shrinkage of the ovary, and assuming this globally reflects a follicular size reduction, we used the apparent ovarian surfaces measured before and after C-Eci treatment (*S1* and *S2* respectively, see Fig. 2D) to calculate a correction factor and estimate the real follicle diameter.

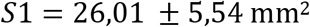

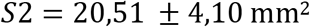

**Figure 3:**
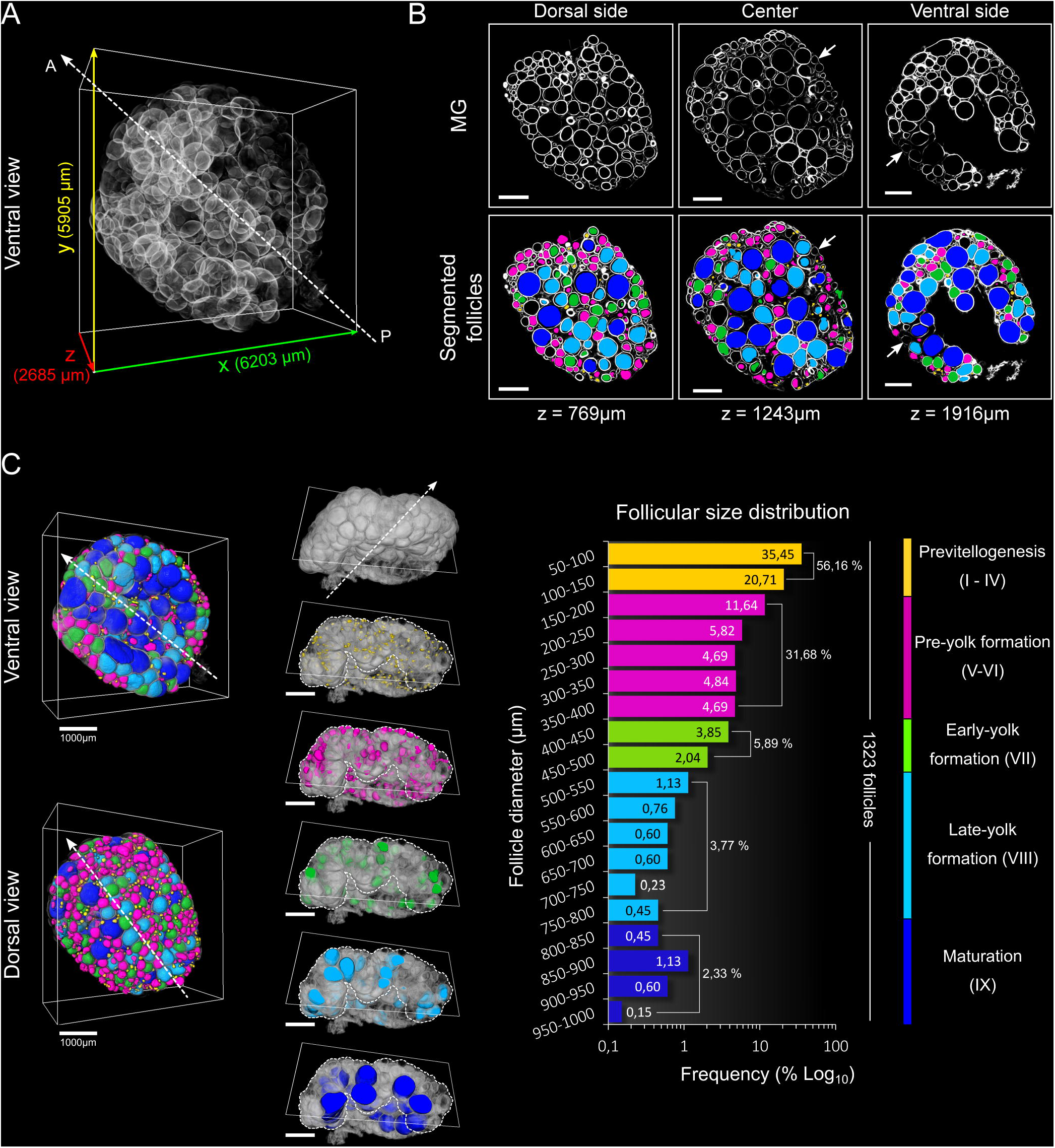
A 3D follicular content analysis in adult medaka ovary. **(A)** 3D reconstruction of a whole medaka ovary after C-Eci clearing, Methyl Green (MG) staining and confocal imaging. **(B)** XY planes at increasing depths (top panels) and the same XY planes merged with follicular 3D segmentation data (bottom panels). Only few follicles are not detected (arrows). **(C)** Ventral and dorsal views of the 3D reconstruction of the whole ovary merged with 3D segmented follicles. Follicles were classified into five classes according to their diameter and the corresponding developmental stage (from stage I to IX). Transversal clipping planes for each of the five classes are shown. White dotted lines delineate the border of the ovary. The large maturing follicles (in dark blue) are preferentially located at the ventral side of the ovary. The follicular size distribution displays the abundance of the small previtellogenic (in yellow) and pre-yolk formation (in magenta) follicles. Frequencies are expressed as a percentage of the total number of follicles. A, anterior; P, posterior. Scale bar: 1 mm.

The correction factor (*corr*) was calculated as follow:

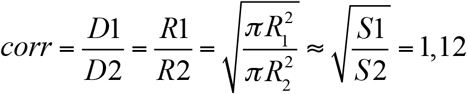

This correction factor was applied to the 1323 follicle diameters values calculated from 3D segmentation data, ranging from 50µm to 963µm, and the follicular size distribution was obtained. Follicles were classified into five categories, corresponding to follicles at different stages of folliculogenesis, from stage I to stage IX (Iwamatsu et al., 1988): previtellogenesis (50-150µm, in yellow), pre-yolk formation (150-400µm, in magenta), early-yolk formation (400-500, in green), late-yolk formation (500-800µm, in light blue) and maturation (>800µm, in dark blue). The resulting follicular size distribution displays a predominance of small previtellogenic (56.16%, in yellow) and pre-yolk formation follicles (31.68%, in magenta). Finally, 3D views of each category of follicles indicate that the large maturing follicles (in dark blue) are preferentially located on the ventral side of the ovary, whereas no evident spatial arrangement could be observed for the other categories.

Using the C-Eci method and confocal imaging, we successfully attempted to image a whole medaka ovary (2685μm thickness) at a cellular resolution and proceeded to semi-automatic segmentation of follicles. It is worth noting that the C-Eci method induces shrinkage of the ovary of about 21% (see Fig. 2D), similarly to the iDISCO+ method that is known to introduce a significant shrinkage of sample estimated at 11% for the mouse brain(Renier et al., 2016). However, a moderate shrinkage may not be considered as a drawback since it can allow easier imaging of large samples such as entire mice(Pan et al., 2016). All the more, we demonstrated that it is possible to calculate a correction factor that can be applied to determine the real follicular diameters.

### 3D imaging of a trout ovary sample after C-Eci and 3D analysis of follicular content

The C-Eci procedure for 3D ovarian imaging was applied to the rainbow trout that has much larger ovaries and oocytes (up to 4-5 mm) than medaka. Pieces of an ovary (about 8×6×2mm) collected from an adult female fish were processed through the C-Eci protocol and MG staining. Opacity was sufficiently reduced to see through the samples (Fig. 4A). A sample was used for 3D imaging by confocal microscopy and the whole volume was reconstructed in 3D (Fig. 4B). No major loss of resolution was observed in XY planes at increasing depths (Fig 4C, left panels). Follicles were segmented from the 3D reconstruction data using a computational semi-automatic 3D segmentation procedure (Fig. 4C, right panels). Follicles located at the border of the volume displayed could not be segmented and few others were not detected, likely due to a discontinuous MG staining (Fig. 4C, arrows). The resulting 3D surface reconstruction of follicles displays different populations of growing follicles of various sizes (Fig. 4D). The same correction factor as for the medaka ovary was used to estimate the real follicular diameters of a total of 499 follicles. The follicular size distribution, ranging from 50µm to 993µm, was obtained. Follicles were classified into 4 categories, corresponding to follicles at different stages from stage2 to stage 5(Estay et al., 2012)(Campbell et al., 2006)(Bromage and Cumaranatunga, 1988): early perinucleolar (50-250µm, in yellow), late perinucleolar (250-450µm, in magenta), cortical alveoli (450-800µm, in light blue) and peripheral yolk granule (>800µm, in dark blue). The resulting profile displays a large predominance of small early perinucleolar follicles (78.36%, in yellow). Finally, no evident spatial arrangement was observed in lamella structures on the 3D view of each category of follicles.

**Figure 4:**
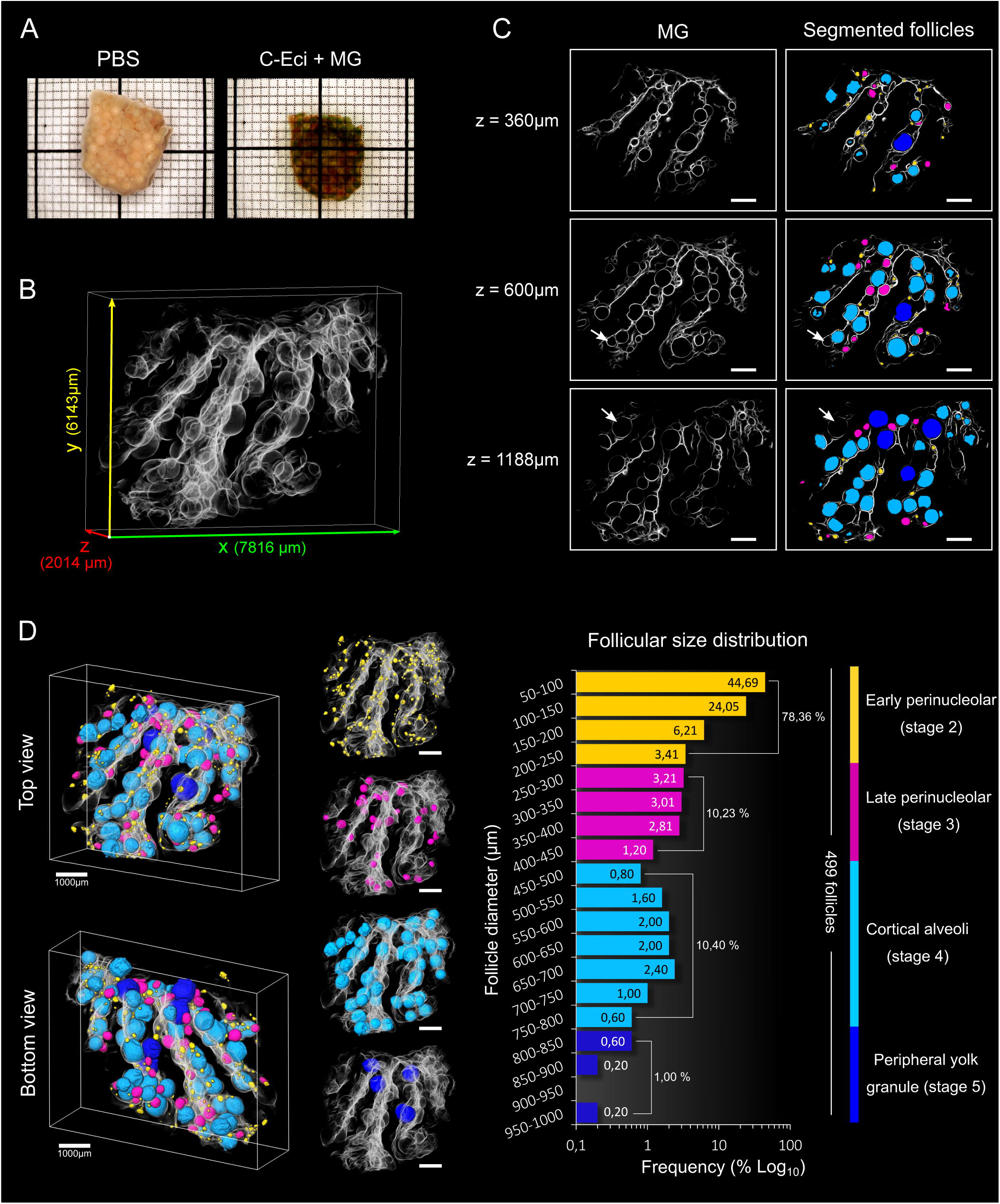
A 3D follicular content analysis in adult trout ovary. **(A)** A piece of trout ovary (2 mm thick) before and after clearing with C-Eci and Methyl Green (MG) staining, showing that C-Eci method efficiently clears trout ovary. **(B)** 3D reconstruction of a piece of trout ovary after C-Eci clearing, MG staining and confocal imaging. **(C)** XY planes at increasing depths (left panels) and the same XY planes merged with follicular 3D segmentation data (right panels). Few follicles are not detected (arrows). **(D)** Top and bottom views of the 3D reconstruction whole sample merged with 3D segmented follicles. Follicles were classified into four classes according to their diameter and the corresponding developmental stages (from early perinucleolar stage to peripheral yolk granule stage). 3D reconstructions for each of the four classes are shown. No evident spatial arrangement of the different class of follicles is observed in lamella structures. The follicular size distribution displays a large predominance of small early perinucleolar follicles (in yellow). Frequencies are expressed as a percentage of the total number of follicles segmented. Square = 1×1 mm; Scale bar: 1 mm.

Using the C-Eci procedure, we could analyze the follicular size distribution in the ovary of both medaka and trout, which was till now limited to late vitellogenic and post-vitellogenic follicles (T, Iwamatsu, 1978). The main limitation of this approach is the minimal size of follicles that could be measured, the tissue shrinkage not being considered as a major drawback as discussed above. The 3D segmentation was indeed restrained to follicles above 50μm in diameter, while 2D segmentation of smaller oocytes could be managed on medaka ovarian sections(Gay et al., 2018). The difficulty with the 3D segmentation of image stacks likely comes from a somehow discontinuous staining contrast in depth. One possibility would be to refine the segmentation algorithm that should greatly benefit from recent machine learning approaches. Anyhow, the C-Eci procedure is a non-toxic and efficient technic for 3D follicular imaging that allows for the first time getting exhaustive and quantitative data on follicles enclosed at a given time in a fish ovary. We now expect that further kinetic analyses of the follicular content throughout the reproductive cycle of both medaka and trout will provide true quantitative data on the oogenesis dynamic in fish ovary. The next step will be to combine the C-Eci protocol with immunostainings for further developmental and physiological analyses, thus opening new perspectives for a better understanding of the biology of the fish ovary.

## MATERIALS AND METHODS

### Ethical statements

All experimental procedures used in this study followed the recommendations of the French and European regulation on animal welfare. Fish rearing and handling were approved by the INRA LPGP-Animal Care and Use Committee (N° Z-2015-30-VT and Z-2015-127-VT-MF) for medaka, and by the INRA PEIMA Institutional Animal Care and Use Ethical Committee for rainbow trout (B29-277-02).

### Fish breeding and sample collection

Adult medaka (*Oryzias latipes*) females from the CAB strain were raised in the INRA-LPGP fish facility at 26°C under an artificial photoperiod (14h light/10h dark), during which the length of the reproductive cycle is of 24h and eggs are daily spawned at the onset of the light. Female rainbow trout *(Oncorhynchus mykiss)* from an autumn-spawning strain were held under natural photoperiod until their first reproduction (2 years) in INRAE-PEIMA experimental fish facilities (Sizun, France). For ovary dissections, medaka (aged from 4 to 5 months) and trout (2.5 years old, 134 days after the first spawning) fishes were euthanized by immersion in a lethal dose of MS-222 solution at 30-50mg/L and 400mg.L^−1^, respectively. Ovary of both medaka and trout were fixed overnight at 4°C by 4% paraformaldehyde (PFA) in 0,01 M phosphate buffer saline pH 7.4 (PBS, Sigma-Aldrich P4417). Ovaries were washed in PBS and conserved in PBS + 0.5% (w/v) sodium azide (S2002, Sigma-Aldrich) at 4°C.

### Optical tissue clearing protocols

#### Simple immersion

All steps were performed at room temperature on an agitator. High-refractive index (hRI) matching solutions were prepared as described in Yang *et al.* with some modifications (Yang et al., 2014a). A 60% sorbitol-based solution was prepared by dissolving 300g of Sorbitol (Sigma-Aldrich S1876) in 400mL of PBS:DMSO (50:50) (v/v). The RI was adjusted at 1.457 by addition of PBS:DMSO. Dilutions of this solution (1:6 and 11:15) were prepared to reach lower RI (1.353 and 1.40). To reach RI 1.49, 120g of sorbitol were dissolved in 50mL of PBS: DMSO. For RI 1.456, 13.32g of Histodenz (Sigma-Aldrich D2158) was dissolved in 10mL of PBS 0.02M (Sigma-Aldrich P4417). For each of the solutions described above, the ovary was immersed successively in 20% (v/v) for 8h; 40% (v/v) for 16h; 60% (v/v) for 8h; 80% (v/v) for 16h and 100% (v/v) for 48h.

#### iDISCO+

iDISCO+ was conducted as described by Renier *et al.* with slight modifications(Renier et al., 2016). All steps were performed at room temperature on an agitator except with dibenzyl ether. Samples were dehydrated in a 20-40-60-80-98% (v/v) series of methanol containing 2% Tween20 for 8 to 16h. A final bath of 100% methanol during 48h was then performed. Ovary was subsequently transferred to 66% dichloromethane (DCM, Sigma-Aldrich 270997)/33% methanol for 3h, and 100% dichloromethane twice. Finally the sample was incubated in dibenzyl ether (DBE, Sigma-Aldrich 108014) for refractive index matching.

#### CUBIC

CUBIC protocol was conducted as described by Suzaki *et al.* with slight modifications(Susaki et al., 2014b). CUBIC-1 reagent is composed by 25% (w/w) urea (Sigma-Aldrich GE17-1319-01), 25% (w/w) N,N,N’,N’-tetrakis(2-hydroxypropyl)ethylenediamine (Sigma-Aldrich 122262) and 15% tritonX-100 (Sigma-Aldrich X-100) in distilled water. CUBIC2 was prepared by mixing 50% sucrose (Sigma-Aldrich S9378), 25% (w/w)urea (Sigma-Aldrich GE17-1319-01), 10% triethanolamine (w/w) (Sigma-Aldrich 90279) and 0.1% TritonX-100 (v/v) in distilled water. Ovary was first incubated in 50%CUBIC-1/50%PBS at room temperature for 3h and then transferred to 100%CUBIC-1 at 37°C on a rotating wheel for three days, with a changing of the solution at 24h. Sample was then rinsed several times in PBS at room temperature and transferred to CUBIC2 for 2 days at 37°C on the rotating wheel.

#### Ethyl cinnamate (ECi)

ECi clearing was adapted from Klingberg et al. with few modifications(Klingberg et al., 2017). All steps were performed at room temperature on an agitator. The sample was dehydrated in a 20-40-60-80-98% (v/v) series of methanol containing 2% Tween20 for 8 to 16h. A final bath of 100% methanol during 48h was then performed and ovary is directly transferred in ECi 100% for a few hours (without shaking). Eci solution is renewed and sample is placed on agitator.

#### CUBIC-1-Eci (C-Eci)

An ovary was first incubated in 50%CUBIC-1/50%PBS at room temperature for 3h and then transferred to 100% CUBIC-1 at 37°C on a rotating wheel for three days, with a changing of the solution at 24h. Samples were rinsed several times in PBS at room temperature and then processed according to the Eci protocol.

### Nuclear staining with Methyl Green

Nuclear staining was performed before clearing steps and after CUBIC-1 in the C-Eci protocol. With these later, the nuclear staining was performed before the serial dehydration steps. Samples were incubated in MG in PBS/0.1% triton (pH7) or in TNT (TrisHCl 0.1M, NaCl 0.15M)/0.1% triton (pH4). MG was used at 80µg/ml or 8µg/ml. Incubations were performed at 4°C or 50°C for 2,5 days. Samples were then washed overnight in PBS + 0.1% tween at room temperature.

### Image acquisition

Bright-field macroscopic images were acquired with an upright Zeiss stereomicroscope with an Axiocam digital camera. For fluorescent imaging, a Leica TCS SP8 laser scanning confocal microscope was used. Samples were imaged with the 16x/0.6 IMM CORR VISIR HC FLUOTAR (Leica) objective. The MG fluorescent signal was detected through laser excitation at 638 nm and fluorescent detection was performed by an internal photomultiplier. Z-steps were fixed at 6 μm for all acquisitions. For clearing method comparisons, samples were imaged with the same settings, including no laser compensation in depth and no image post-processing. For 3D whole sample imaging, images were acquired with laser compensation and contrast enhancement was applied. For medaka ovary and because of its large size (>2,5 mm), the dorsal and ventral parts were imaged separately. For each face, a total volume of about 7.5 x 7.5 x 2 mm was acquired (voxel size: 1.8 x 1.8 x 6 μm) and generated 5GB of data. The overlap was of 1 mm. All samples were imaged within the following week after staining and clearing.

### 3D image reconstruction and segmentation

Whole specimens imaged by confocal microscopy were treated with the Amira 2019.2 software (Thermo Fisher Scientific) on a 64-bit Windows 10 Pro computer equipped with a 2x Intel Xeon Silver 4110 (8 Core, 3.0GHz) processor, a Nvidia Geforce GTX 1080 graphic card and 384 Go of RAM. To minimize the data volume, raw data initial size (spatial resolution 1,8 x 1,8 x 6 µm) were scaled to 0.3 (1:3) in x and y. 3D segmentation of follicles was performed semi-automatically in the project editor of Amira using a combination of different operation with systematic 3D interpretation. To binarize the nuclear signal, the image gradient (canny deriche) was calculated and a Top-Hat was applied on resulting images to threshold nuclei. Images were inverted to visualize the internal part of follicles and connective tissues. Morphological operator like opening and hole fill was performed to improve follicles shapes. A watershed was applied to separate connected object. To eliminate non-follicles remaining structures, binary objects were filtered based on their size and shape. Only follicles with a spherical shape and a size above 50µm^3^ were kept. Resulting 3D reconstruction of follicles was generated with Amira’s “Volume rendering” visualization module. Volume of all segmented follicles were exported, equivalent diameters were calculated and corrected with a correction factor to compensate the volume shrinkage due to sample clearing. Follicles were then colorized by diameter range, based on their corrected size.

### Statistics

Statistical analyses were performed by using Rstudio Version 1.1.463. P-values were calculated by using a non-parametric Wilcoxon signed-rank test for indicating significant differences between samples.

## ACKNOWLEDGMENTS

We thank the INRA-LPGP fish facility staff, and especially Cecile Duret, for fish rearing and husbandry.

## AUTHOR CONTRIBUTIONS

ML performed image processing, data analyses and participates in manuscript writing. MT performed samples clarification, samples imaging, led the set up of the C-Eci protocol and participates to the manuscript writing. JBu performed the semi-automatic 3D segmentation workflow. AB participated to the set up of the clarification protocol. SG and EC collected samples. JB participated in the design of the study. VT conceived the study, participated in data analyses and manuscript writing. All authors read and approved the final manuscript.

